# Quasi-Entropy Closure: A Fast and Reliable Approach to Close the Moment Equations of the Chemical Master Equation

**DOI:** 10.1101/2021.12.01.470753

**Authors:** Vincent Wagner, Benjamin Castellaz, Marco Oesting, Nicole Radde

## Abstract

**Motivation:** The Chemical Master Equation is the most comprehensive stochastic approach to describe the evolution of a (bio-)chemical reaction system. Its solution is a time-dependent probability distribution on all possible configurations of the system. As the number of possible configurations is typically very large, the Master Equation is often practically unsolvable. The Method of Moments reduces the system to the evolution of a few moments of this distribution, which are described by a system of ordinary differential equations. Those equations are not closed, since lower order moments generally depend on higher order moments. Various closure schemes have been suggested to solve this problem, with different advantages and limitations. Two major problems with these approaches are first that they are open loop systems, which can diverge from the true solution, and second, some of them are computationally expensive.

**Results:** Here we introduce Quasi-Entropy Closure, a moment closure scheme for the Method of Moments which estimates higher order moments by reconstructing the distribution that minimizes the distance to a uniform distribution subject to lower order moment constraints. Quasi-Entropy closure is similar to Zero-Information closure, which maximizes the information entropy. Results show that both approaches outperform truncation schemes. Moreover, Quasi-Entropy Closure is computationally much faster than Zero-Information Closure. Finally, our scheme includes a plausibility check for the existence of a distribution satisfying a given set of moments on the feasible set of configurations. Results are evaluated on different benchmark problems.

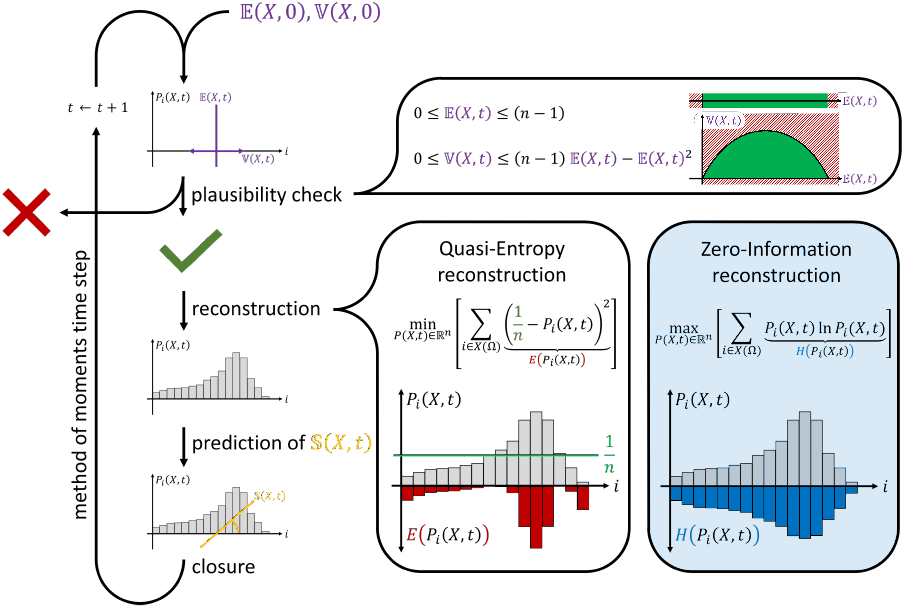

## Introduction

Many intracellular processes such as signal transduction pathways, gene regulatory networks or metabolic processes are described as (bio)chemical reaction networks. In recent years, stochastic modeling approaches have become more and more frequently used in connection with single cell data to capture heterogeneities arising from the stochastic nature of reaction events (1). The Chemical Master Equation (CME) is a set of linear differential equations which describes the time evolution of the probability distribution over all possible configurations of the system. Unfortunately, it is usually computationally not tractable to solve the CME directly, since the number of configurations is huge even for small systems, leading to high-dimensional differential equation systems that cannot effectively be stored or solved. A possible remedy is the Stochastic Simulation Algorithm (SSA), which generates sample paths from the CME (2, 3). However, generating sufficiently many sample paths to properly capture the system’s behavior is computationally expensive which also renders this method sub-optimal.

A very elegant approach for calculating the main characteristics of the solution of the CME is called the Method of Moments (MoM). Based on the CME, it derives a set of coupled Ordinary Differential Equations (ODEs) which describe the time evolution of the statistical moments of the CME solution (4). Assuming that the solution of the CME can be described by a few moments reduces the computational complexity drastically, since the dimension of the ODE system reduces from the number of configurations to the number of moments. If the reaction system only contains reactions of order zero or one, the moment equations are closed and can directly be integrated. However, for chemical systems involving propensities of order two or higher, one key challenge when solving the ODE system arising from the MoM is that moments of order *j* depend on higher order moments. In other words, the ODE system is not closed and can therefore not be solved directly.

Several moment closure schemes exist. The most common and trivial approach is to simply truncate the equation system by setting all higher order moments to zero. We refer to this closure as Truncation Closure (TC). It is easy to implement but relies on the assumption that the influence of higher order moments on the regarded moments can be neglected. This assumption is oftentimes violated (5, 6). Besides TC, several other types of closure scheme have been suggested. Many approaches assume that the solution of the CME is similar to a distribution of known form or a member of a specific distribution family (see e.g. Whittle (7), Krishnarajah et al. (8, 9), Ramalho et al. (10), Lakatos et al. (11), Bronstein and Koeppl (12)) and construct the closing moments accordingly. Singh and Hespanha (13, 14, 15) derived and extended a derivative-matching technique. Keeling (5) replaced the addition of higher order moments with a multiplicative term and subsequently closes the system on this basis. The studies of Soltani et al. (16) and Naghnaeian and Vecchio (17) suggest to derive higher-order moments as nonlinear functions of (conditional) lower order moments. Ruess et al. (18) couple the MoM solution to samples of the true CME solution with the help of a Kalman filter on SSA trajectories. Hasenauer et al. (19) and Kazeroonian et al. (20) divide all involved species into groups with low or high corresponding copy numbers, respectively. While species with low copy numbers are modeled as discrete stochastic quantities, statistical moments conditioned on the abundance of low copy number species are used to describe the abundance of all other species.

In addition, we want to discuss one method in more detail. The Zero-Information Closure (ZIC) introduced by Smadbeck and Kaznessis (21) reconstructs a distribution in every time step to subsequently calculate higher order moments from it. To this end, a constrained optimization problem is solved that maximizes the information entropy while still returning a distribution possessing the first *J* given moments. The approach takes roots in the idea of introducing as little artificial information as possible into the solution of the MoM ODE. Another advantage is that ZIC produces moment trajectories and corresponding distributions simultaneously. On the downside, the method is computationally expensive and oftentimes numerically unstable, leading to diverging solutions even for simplistic systems. Andreychenko et al. (22) circumvent the runtime issues, as they only reconstruct a corresponding distribution at the very final time step. However, this often comes at the expense of a low prediction accuracy, since they use TC for the integration of the ODE.

While some methods are more involved than others, one unifying aspect of all published approaches is that there is no a priori result specifying the number of moments required to achieve a certain solution quality for general systems. Solutions can in fact significantly differ from the true moments depending on the system under investigation, the parametrization and the number of moments which are considered (6). For the special case of monostable systems, Grima (23) compares TC to the system size expansion of van Kampen (24) and concludes that TC delivers meaningful results in this context.

Beyond this specific setting, Schnoerr et al. (6) generally discuss under which conditions TC returns physically meaningful results. In this context, a physically meaningful result needs to fulfill four properties: (i) Mean molecule numbers and all central moments of even order are positive, (ii) moment trajectories converge to a steady state whenever the CME itself has a stationary solution, (iii) the moments approach a unique steady state independent of the initial conditions, and (iv) moments do not oscillate. On this basis they conclude that physically meaningful moment trajectories can be reached for certain systems and parameter ranges. In a consecutive work, Schnoerr et al. (25) compare the results of different aforementioned closure schemes with a focus on physical meaningfulness and conclude that meaningful parameter ranges for general systems are largest when using TC. Unfortunately, they restrict themselves to systems with irreducible configuration spaces and do not present plausibility criteria beyond non-negativity of mean molecule numbers and even order central moments.

Our contribution to the presented challenges is two-fold: First, we introduce the Quasi-Entropy Closure (QEC), a variant of ZIC, which closes the moment equations by minimizing the distance to the uniform distribution subject to the first *J* moments as constraints. This leads to a quadratic objective function, which can be solved much more efficiently and stably compared to entropy maximization by standard optimization algorithms. To assess the simulation quality, we complement QEC with a set of powerful and mathematically necessary conditions that serve as a plausibility check, indicating impossible moment trajectories on the fly. These plausibility criteria use the dependencies between moments to restrict higher order moments for distributions on a bounded space of configurations.

We demonstrate on different example systems that QEC out-performs ZIC in terms of computation time for single distribution reconstruction steps and overall simulation time. Moreover, we show that QEC still gives good results if the number of optimization steps in each reconstruction step is significantly lowered, which is not the case for ZIC. Finally, we discuss results on a system for which a closure of two moments fails. In this case, the plausibility check is able to detect the resulting divergence from the true underlying solution and suggests to increase the number of moments, which solves the problem so that final results are in good agreement with the true underlying solution.

## Systems and Methods

### Systems

We apply QEC to three systems that prototypically represent a large variety of biochemical systems (Figure 1). For brevity of notation and ease of understanding, we consider systems whose configuration space can be reduced to one single species. Here, the configuration *X* of the system corresponds to the number of *A* molecules in all examples.

**Fig. 1.**
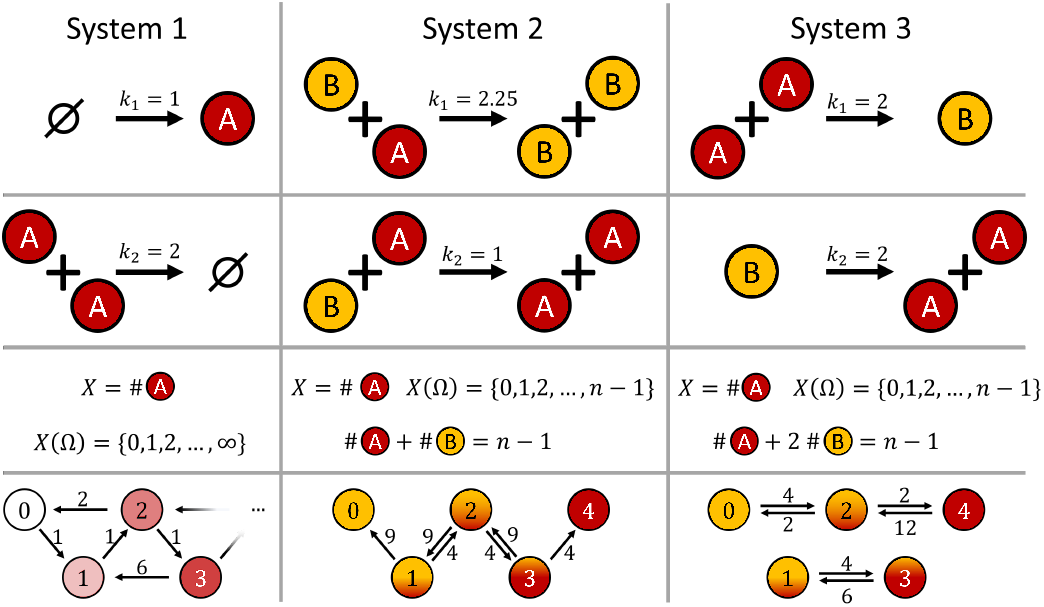
Chemical reaction systems considered in this study. Reaction schemes, space of configurations and Markov transition graphs for the three systems featured in this study. All reaction rate constants are assumed to be strictly positive.

### System 1

System 1 consists of a constant production rate and quadratic degradation and is therefore one of the most simple systems involving quadratic propensities:

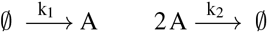

Due to the irreducible configuration transition graph, it converges to an equilibrium distribution. Furthermore, this system has an infinite configuration space *X*(Ω).

### System 2

This system is characterized by two reactions with the same reactants *A* + *B* that either react to 2*A* or 2*B*:

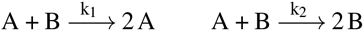

This system has a finite configuration space *X*(Ω) = {0, 1, …, *n* − 1} and the transition graph has two absorbing system configurations, *X* = 0 and *X* = *n* − 1, where *n* − 1 is the total number of molecules in the system. Steady state distributions are given by

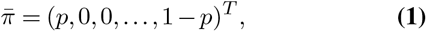

where the value of *p* ∈ [0, 1], which is the probability to reach the absorbing configuration *X* = 0, depends on the initial configuration of the system. Thus, the Markov process does not converge in a Markovian sense.

### System 3

System 3 represents a reversible dimerization reaction:

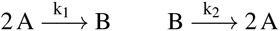

The configuration transition graph consists of two separated subgraphs, one for an even number of *A* molecules and one for an odd number. Each subgraph is irreducible and the Markov process converges to an equilibrium distribution on this subgraph. Moreover, System 3 also has a finite configuration space *X*(Ω) = {0, 1, …, *n* − 1}.

## Methods

The CME describes the time evolution of the probability *P* (*X* = *i, t*) =: *P*_*i*_(*X, t*) that the system is in configuration *X* = *i* at time *t* and is given by

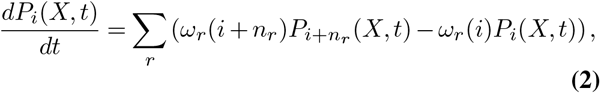

where *ω*_*r*_(*i*) is the propensity for the *r*-th reaction conditional on being in configuration *i* and *n*_*r*_ is the corresponding configuration change vector. The solution of Eq. (2) is a time-dependent probability distribution *P* (*X, t*) on the discrete space of configurations *X*(Ω).

The MoM computes the first *J* moments of this distribution, i.e. the expected value

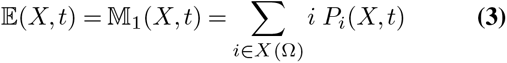

and the centered moments of order *j*

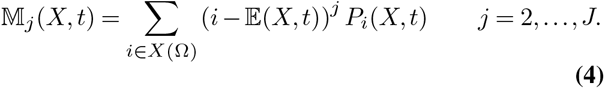

Evolution equations for the moments can be derived directly from the CME (4). Assuming that all propensities are polynomials in *i*, the resulting equations have the form

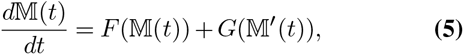

where 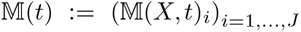 denotes the vector containing the first *J* moments and 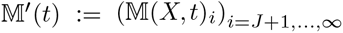 denotes the sequence containing all moments of higher order (21). *F* and *G* are polynomials in 𝕄 and 𝕄 ′, respectively.

If all propensities are at most linear, i.e. of order 0 or 1, then *G ≡* 0 and Eq. (5) is a closed system of ODEs. If at least one propensity has order two or higher, *G ≢* 0 and Eq. (5) is no longer a closed system of ODEs. Closure schemes are approximations for 𝕄 ′ in order to close Eq. (5). Moment equations for Systems 1,2 and 3 are specified in Supplementary Note 1.

An interesting fact, which to the best of our knowledge has only been considered by Smadbeck and Kaznessis (21) so far, is that, given 𝕄, the values of 𝕄 ′ are restricted to a certain range. Consider a system with *n >J* different configurations. Then, Eq. (3) and Eq. (4) together with the conditions that

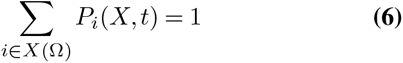

and

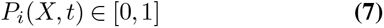

define a manifold for *P* (*X, t*). The range of reasonable values for 𝕄 ′ is restricted by this manifold. The interesting question now is, how does one choose a reasonable distribution on this manifold? Smadbeck proposed the appealing approach of choosing the distribution that maximizes Shannon entropy (21). If one has no further information about a system, this choice seems reasonable because it maximizes uncertainty. However, this approach has the disadvantage that the resulting optimization problem is computationally very hard in the number *n* of configurations – the problem that the MoM originally intended to avoid.

Our method is based on a similar argument to Smadbeck’s approach, but instead solves a substitute problem that is computationally easier (and much more stable) to solve. Instead of maximizing the entropy of the distribution, we minimize the mean squared distance to the uniform distribution. If no moment constraints are considered, both optimization problems have the same solution, a uniform distribution on the discrete configuration space.

## Algorithm

Our approach is illustrated in Figure 2. The system is initialized for *t* = 0 by providing the first *J* moments of the initial distribution, e.g. the expectation 𝔼 (*X, t* = 0) and variance 𝕍 (*X, t* = 0) for *J* = 2. Then, a plausibility check is performed in each time step before the next integration step in order to detect unfeasible solutions, as explained below. If solutions are feasible, the algorithm proceeds by closing the set of differential equations for the moments. This is done via reconstructing the distribution which obeys the first *J* moments at the current time *t* and has minimal distance to a uniform distribution. To this end, a constrained optimization problem is solved, as outlined below.

**Fig. 2.**
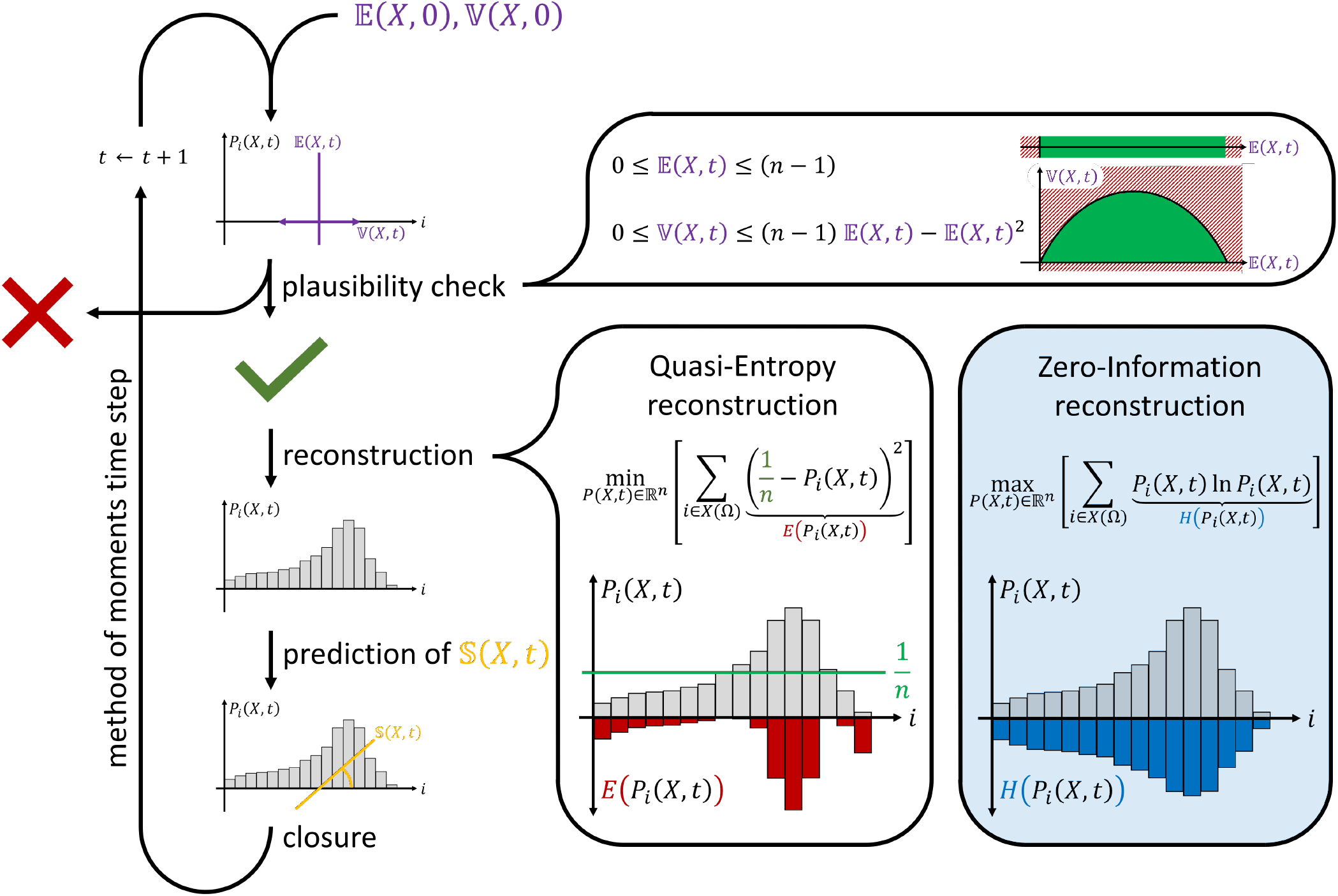
Schematic representation of QEC for the MoM. The system is initialized at *t* = 0 by the first *J* moments of the initial distribution *P* (*X, t* = 0). At each time instance, universal inequalities for the moments of a distribution on a bounded configuration space are used to check whether the proposed moments are feasible, i.e. whether there exists a distribution with those moments. If this is not the case, this is a clear indication that the moment equations deviate from the true underlying solution, and simulation quality has to be improved. If the plausibility check is positive, the moment equations are closed by reconstruction. In this step, an optimization problem is solved which searches for the distribution that minimizes the configuration-wise quadratic distance *E*(*P*_*i*_(*X, t*)) to the uniform distribution constraint to the given moments. This distribution is then used to calculate higher order moments which are needed to close the moment equations. Smadbeck and Kaznessis (21) use a similar approach for the reconstruction, in which the information entropy is maximized during the reconstruction step.

After reconstructing the distribution, higher order moments are calculated and used for the next integration step of the moment equations. For *J* = 2 and a system with reactions up to second order, for example, the skewness 𝕊 (*X, t*) is sufficient to close the equation system.

### Reconstruction step

Reconstruction is achieved by solving an optimization problem for each time instance *t*,

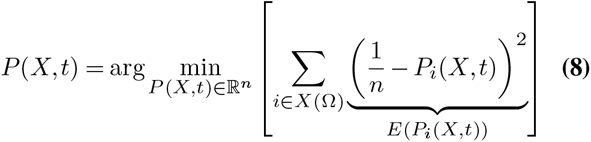

subject to

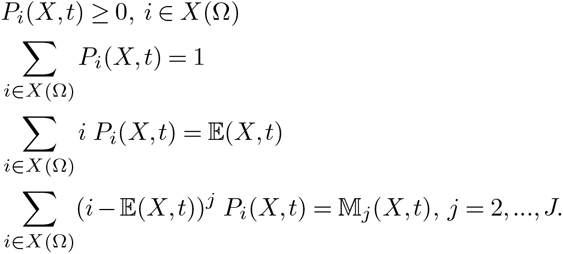

i.e. we search for the distribution which is maximally close to a uniform distribution on the configuration space *X*(Ω) constraint to having the *J* given moments. This problem can be reformulated with Lagrange parameters *λ*_0_, …, *λ*_*J*_, giving as a solution (see Supplementary Note 2)

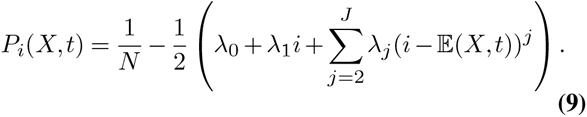

In contrast, ZIC reconstructs the distribution by maximizing the information entropy,

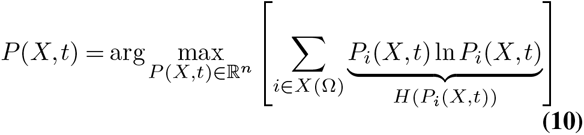

as indicated in the blue box in Figure 2.

Both optimization problems are convex. If moments are introduced as constraints, we anticipate that the QEC optimization problem Eq. (8) is easier to optimize mainly due to two reasons. First, in order to solve a convex optimization problem, one searches for the roots of the gradient of the objective function. In case of information entropy, this gradient becomes large for small probabilities, making gradient-based approaches inherently unstable. Second, a typical applicant of chemical simulation methods will oftentimes resort to preimplemented solvers for such optimization problems. The `python` standard library `scipy`, for example, incorporates two solvers capable of solving such problems. While the sequential least squares programming solver is intrinsically designed to solve quadratic problems, the trust-region algorithm ensures constraint satisfaction by estimating a trust region based on a quadratic representation of the true error function. In both cases, the problem structure proposed by us is advantageous to the original formulation.

### Plausibility check

Our plausibility check, which is applied in each time step, provides sufficient conditions to detect unfeasible moment trajectories. It uses dependencies between moments of different orders and is valid for any probability distribution *P* (*X*) on a bounded configuration space *X*(Ω) = {0, 1, …, *n* – 1}. Bounds for a moment of order *j* + 1 are calculated using all lower order moments in the following way:

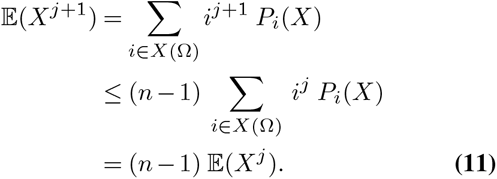

For the variance, this gives

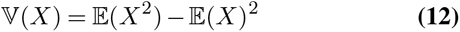

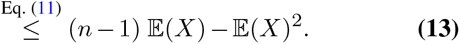

The skewness can be bound in a similar fashion:

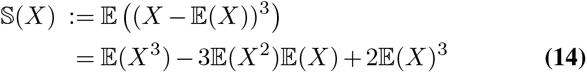

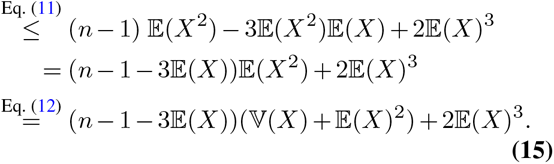

While the expected value 𝔼 (*X*) and all centered higher order moments of even order have a trivial lower bound of zero, this is not the case for centered moments of odd orders larger than one. Due to symmetry reasons, however, Eq. (15) should generally hold for |𝕤| and not only 𝕤.

This derivation procedure can be applied to centered moments of arbitrary order, even though the resulting inequalities become extensive in length. A bound for the curtosis is presented in Supplementary Note 3A. We note that the inequalities Eq. (13) and Eq. (15) are sharp, i.e. there exist distributions for which the inequalities become equalities, as outlined in Supplementary Note 3B.

These inequalities are an efficient way to assess the solution quality on the fly. Before each closure step, they are used to monitor whether a distribution on the finite configuration space with those moments can exist. If the moments fail to fulfill the moment inequalities, the proposed moment trajectories cannot be meaningful. Consequently, this indicates that the simulation quality needs to be improved, as we will discuss in more detail below.

### Implementation

#### Simulation settings

All results presented in this work are generated using the same `python 3` code framework (26). We implemented the MoM ODE system on the basis of the python standard libraries `numpy` (27) and `scipy` (28) and used `matplotlib` (29) to generate graphics. The time integration is performed using the standard solver `RK45`. The simulation time was set to 1*s* for all virtual experiments.

While TC is trivial to implement, the optimization problems arising from ZIC and QEC are solved using the `SLSQP`-Optimizer from `scipy`. We also tried the only other alter-native `trust-constr`, but did not use it for further studies due to worse overall performance. Each optimization problem was initiated with the discrete uniform distribution over the space of configurations. We used the `SLSQP`-Optimizer with either 250 maximally allowed optimization steps or with only 25 steps for a reduced runtime, as indicated in the figure captions.

In contrast to Smadbeck and Kaznessis (21), our MoM results are compared to the solution of the CME itself instead of SSA simulations. For System 1, a configuration space truncation is needed so that the resulting CME is finite-dimensional. A truncation to *n* = 21 configurations in combination with our chosen parameter sets (see Figure 1) ensures that the sum over the solution of the CME approaches 1 up to machine precision for all regarded time steps. For both other systems we also used *n* = 21.

#### QEC outperforms ZIC in terms of runtime

Figure 3 shows the performance of QEC in comparison to TC and ZIC for System 1 and the MoM for the first three moments 𝔼, 𝕍 and 𝕤. Figure 3A shows inferred moment trajectories. The reference solution is shown in yellow and corresponds to the moments directly obtained by solving the CME numerically. This comparison is different to that in Smadbeck and Kaznessis (21), who used SSA simulations as reference. For ZIC and QEC, each dot on the line corresponds to one distribution reconstruction step. ZIC and QEC both give accurate solutions which are close to the reference trajectories for the entire integration range, while TC deviates from the reference values for all moments. This deviation is still acceptable for the expectation value, however, the error becomes larger for variance and skewness. The frequencies of the computation times for single reconstruction steps for QEC and ZIC are depicted in Figure 3B. This diagram clearly shows that, due to the nature of the optimization problem, the reconstruction time is much shorter for QEC compared to ZIC for all reconstruction steps. The average difference is about two orders of magnitude, which means that a single reconstruction is on average about 100 times faster in QEC compared to ZIC. While TC solely works with moments and does not allow for a reconstruction of the underlying distributions, this is possible for ZIC and QEC. A continuous interpolation of reconstructed distributions is shown in Figure 3C for ZIC (blue) and QEC (red) in comparison with the reference CME solution (yellow). The course of distributions inferred by ZIC and QEC are quite similar, and both do not capture the multimodal nature of the transient behavior of the reference solution. However, the steady state distribution is accurately captured by both methods.

**Fig. 3.**
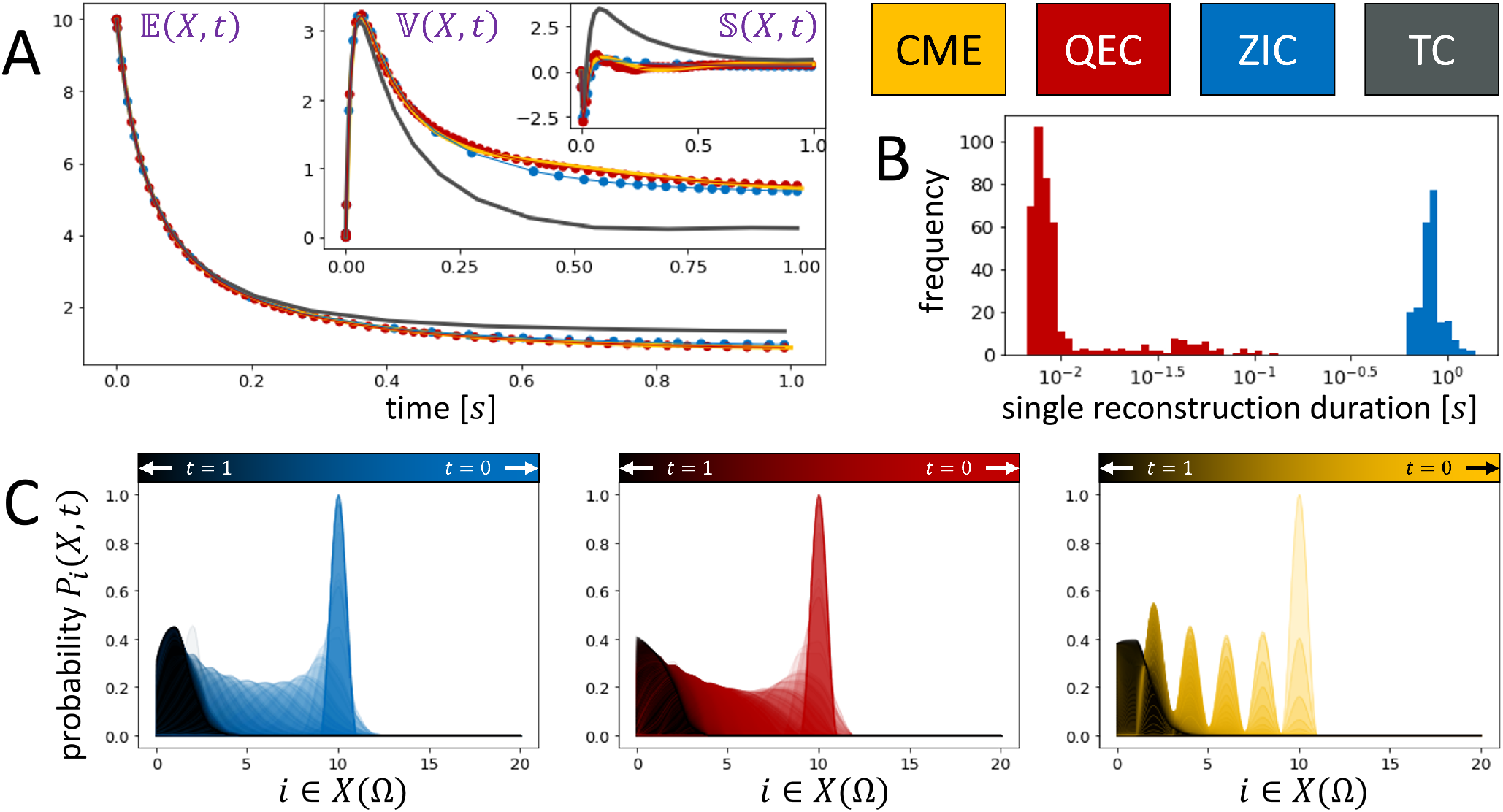
QEC and ZIC outperform TC and QEC has much faster runtime. Comparison of MoM results for different closure schemes for System 1 with the first three moments and *X*(*t* = 0) = 10 and *n* = 21. Reference moment trajectories and distribution courses obtained from the CME are depicted in yellow. For ZIC and QEC, a maximum of 250 optimization steps was allowed per distribution reconstruction. (A) Courses of 𝔼, 𝕍 and 𝕤 for ZIC (blue), QEC (red) and TC (gray). For ZIC and QEC, each dot on the line corresponds to one distribution reconstruction step. (B) Histogram of computation times for single reconstruction steps. (C) Continuous interpolation of the reconstructed discrete distributions for ZIC and QEC in comparison to the reference course. Light colors represent early time steps, darker colors correspond to reconstructions in later steps.

Summarizing, these results show that ZIC and QEC both outperform TC on this example in terms of accuracy of the moment trajectories, and that calculated moments of both methods are close to their reference values. While inferred distributions are similar among both methods, reconstruction is much faster for QEC. Results furthermore demonstrate that three moments for this system are not sufficient to capture the transient course of the true underlying distribution, but give satisfactory results for the steady state distribution.

Results for Systems 2 and 3 show very similar behavior and can be found in Supplementary Note 4, Figures 6 and 7. Also here, TC moments deviate from the reference trajectories for both systems, while ZIC and QEC give accurate results for the moment courses. Reconstructed distributions are similar for both methods and in case of System 2 also capture the course of the reference distribution, while this is not the case for System 3, which also has a multi-modal behavior.

#### QEC is able to reconstruct moments using a reduced number of optimization steps

We asked the question whether the overall runtime for ZIC and QEC could be reduced by lowering the number of optimization steps in each reconstruction step. Therefore, we reduced the maximum number of optimization steps from 250 to 25 for ZIC and QEC (Figure 4). Moment courses (Figure 4A) show that ZIC fails completely with this low number of optimization steps, while the results of QEC are similar to those of Figure 3, which have been obtained with a maximum of 250 optimization steps in each reconstruction. Moreover, computation times (Figure 4B) show that QEC is still more than one order of magnitude faster for a single reconstruction step in this setting. The failure in the moments is also visible in the course of reconstructed distributions (Figure 4C), which indicates that ZIC indeed needs a much larger number of optimization steps to converge to an optimum.

**Fig. 4.**
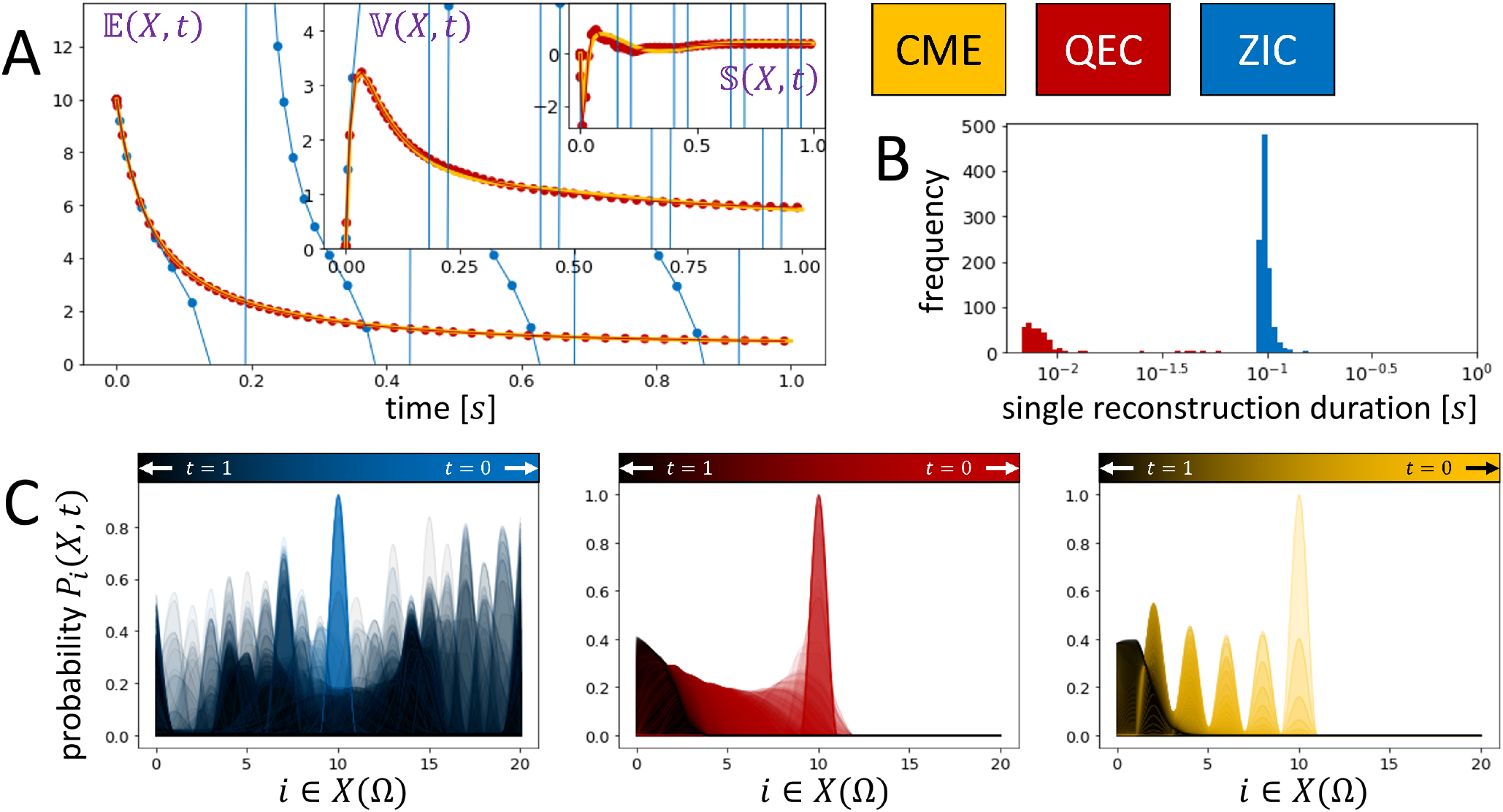
QEC only requires a low number of maximal optimization steps compared to ZIC. Results similar to Figure 3 with the maximal number of optimization steps set to 25 for QEC and ZIC for each reconstruction step.

Overall, these results show that QEC needs much less optimization steps to infer accurate moment trajectories than ZIC. Thus, the overall runtime of QEC can further be reduced by optimizing the number of iterations in each reconstruction step.

Results for Systems 2 and 3 are shown in Supplementary Note 4, Figures 8 and 9. Again, ZIC fails completely for both systems in this analysis, demonstrating that maximization of the information entropy is much more challenging from an algorithmic viewpoint and 25 optimization steps are not enough to find a good solution. This is different for QEC, which gives good results in terms of moments. As expected, reconstructed distributions are again similar to the reference for System 2, but not for System 3.

#### Plausibility criterion as a quality check to improve MoM results

The results of the plausibility check can be used to assess the feasibility of the inferred moments on the fly. It can be included into the workflow procedure by increasing the number of moments as soon as the criteria are not fulfilled any more at a particular time instance after an integration step, as shown in Figure 5A. For example, when the initial number of moments for the MoM are set to two, *J* = 2, and the plausibility check fails at a time instance for either the expectation value or the variance, the calculations are stopped and QEC is re-initialized with three moments. This procedure is repeated until moment trajectories are reconstructed without any failures of plausibility checks. This is exemplarily shown for System 2, which naturally has a bounded configuration space and therefore allows to evaluate the inequalities for the plausibility check without any further argumentation (Figure 5B). Reconstruction with *J* = 2 moments fails, and moment trajectories deviate from the reference solutions already at an early time point. This is eventually detected by our plausibility criteria in the course of the variance at about *t* = 0.375*s*, where the variance trajectory violates the upper bound condition. Computations with *J* = 3 moments are sufficient for an accurate reconstruction.

**Fig. 5.**
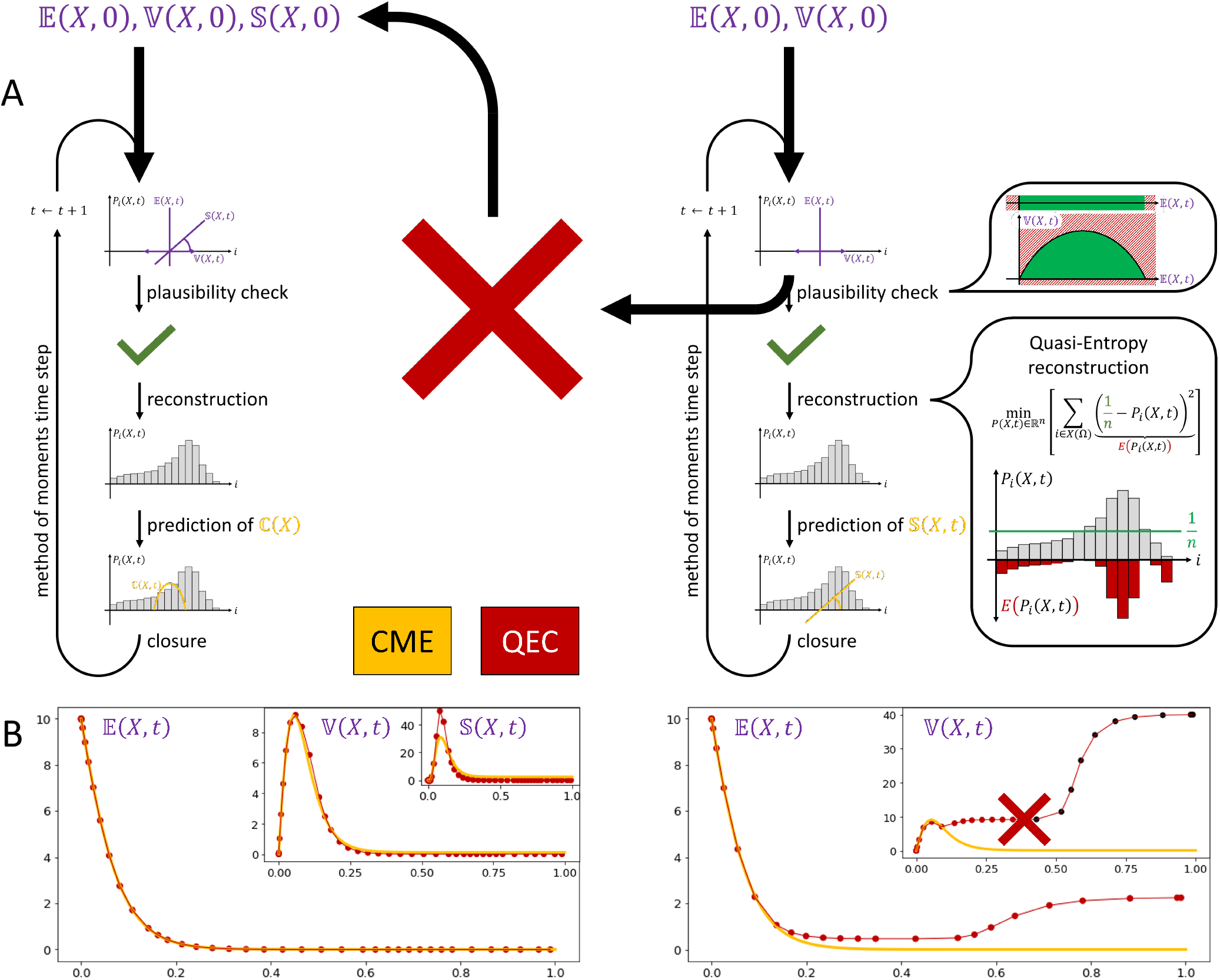
Inclusion of plausibility check into QEC scheme allows to detect deviations from reference trajectories. The plausibility check tests after each integration step whether necessary inequality conditions for the calculated moments are fulfilled. If this is not the case for any of the moments, calculations are stopped and the procedure is re-initialized with an increased number of moments. This is repeated until no violations are detected any more. (A) General procedure, (B) Approach applied to System 2 with *J* = 2 moments, *X*(*t* = 0) = 10 and *n* = 21. The variance deviates from the reference solution already at an early time point, which is eventually detected by the plausibility check, as indicated by dark points. QEC is stopped and re-run with *J* = 3 moments, which is sufficient here to give accurate results.

## Discussion

In this work, we have introduced QEC as a closure scheme which estimates higher order moments of the solution of the CME by finding the distribution which minimizes the distance to a uniform distribution constrained to obey the first *J* moments. Our approach was inspired by ZIC, which reconstructs the optimal solution by maximizing the information entropy under the same moment constraints. We discuss the differences that are implied by this change in objective function from an optimization point of view and demonstrate on three benchmark example systems that QEC is computationally much more efficient than ZIC. Moreover, our approach integrates a plausibility check which is based on bounds for higher order moments depending on lower order moments. These inequalities are generally valid for bounded configuration spaces. Our results show that this plausibility check is able to detect a significant deviation of the calculated moments from the reference moment trajectories, and the QEC scheme is able to react accordingly to improve accuracy of the solution. This can for example be done by increasing the number of moments for the closure scheme.

Interestingly, we observed that the integration step size, which determines how many reconstruction steps are performed and therefore impacts total computation times, varies considerably across the three benchmark examples, and also between ZIC and QEC.

A crucial assumption for the MoM to work properly is that the time dependent distribution over the configurations of the system can accurately be described by a few moments for each time instance. This is often the case, but we do not have reliable a priori estimates when this is the case and how many moments are required. This is a general drawback inherent to all moment closure schemes and the reason why it often fails to predict the true system behavior adequately (5, 6). It is for instance known that capturing multi-modality generally requires more moments than unimodal distributions (19). The shortcomings of the MoM are based on its open loop construction: Except for the initial time instance, there is no coupling between the ODEs derived from the MoM and the actual physics of the underlying system. In particular, the ODE system does not incorporate basic properties such as non-negativity of the expected value and centered moments of even order. The necessary conditions derived as moment inequalities in our plausibility check do not solve this problem completely, but we see it as a step towards detecting deviations from the reference moments at an early stage of the computations. However, it only provides sufficient conditions. This certainly leaves room for further development in the future.

As the moment inequalities are based on a bounded configuration space, they are not directly applicable to systems with an infinite or unbounded space of configurations, as System 1. In practice, however, one can often truncate the configuration space anyway by only considering those configurations which have a probability above a certain threshold.

An advantage of closure methods such as ZIC and QEC compared to TC is that they allow to reconstruct the entire distribution and not only its moments, which sometimes can be used as an additional quality check if anything about the distribution is known. On the other hand, the reconstruction of a distribution makes those methods computationally demanding, since it requires the estimation of probabilities for each configuration of the system at each time instance. Thus, QEC, as any other closure scheme with a reconstruction step, scales very poorly with increasing molecule numbers, as this increase in molecule numbers typically expands the number of configurations tremendously. QEC is thus so far only applicable to small benchmark systems with a few reactions and a relatively small number of molecules.

Another issue with the MoM is that the ODE system can become stiff as higher moments enter the computation, since the moments are likely to differ by several orders of magnitude. This requires solvers which can cope with stiffness. Moreover, when using moment equations for an inverse model analysis, e.g. in parameter estimation, this might require adaptations, as e.g. discussed in Lück and Wolf (30).

In conclusion, we built on the well-founded ZIC and improved it with regard to simulation speed and numerical stability, thereby broadly expanding its application possibilities. Our plausibility criteria complement the general concept of the MoM with a powerful set of necessary conditions that determine physically meaningless simulation trajectories on the fly, consequently avoiding unnecessary simulations.

## Abbreviations

(CME): Chemical Master Equation
(MoM): Method of Moments
(ODE): Ordinary Differential Equation
(QEC): Quasi-Entropy Closure
(SSA): Stochastic Simulation Algorithm
(TC): Truncation Closure
(ZIC): Zero-Information Closure

## Data availability

Our code and files are accessible via FAIRdom Hub (https://fairdomhub.org/models/798?version=1) and can be executed with any common `python 3` distribution.

## Acknowledgments

This preprint was formatted using a L^A^TEX class by Ricardo Henriques that can be accessed here.

## Supplementary Note 1: Moment Equations for Systems 1, 2 and 3

The moment equations for System 1 read

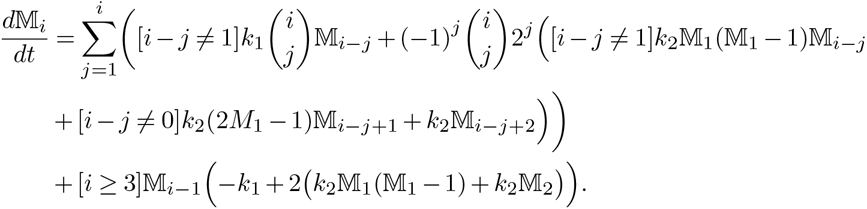

The ODE system for System 2 is given by

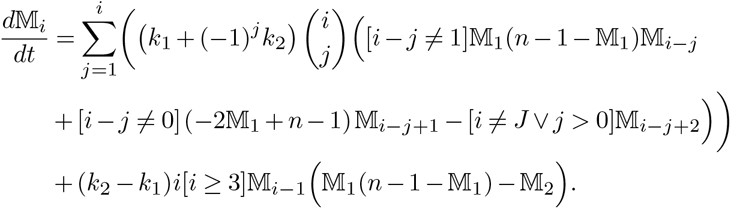

For System 3, we obtain

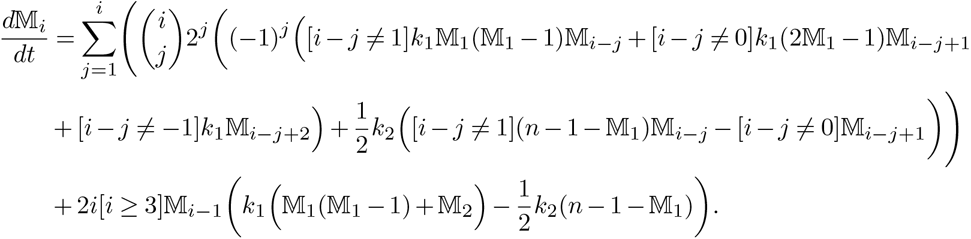

## Supplementary Note 2: Solution of Constrained Optimization Problem via Lagrange Multipliers

The optimization problem of QEC,

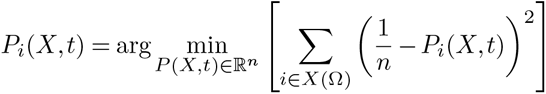

subject to

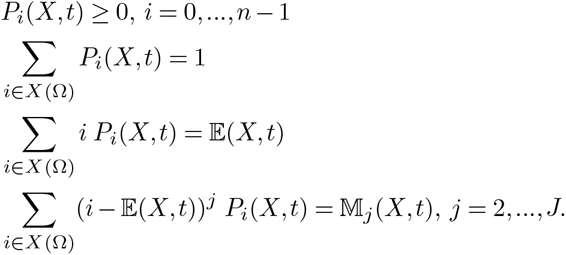

can, at least for the equality constraints, be solved via Lagrange multipliers:

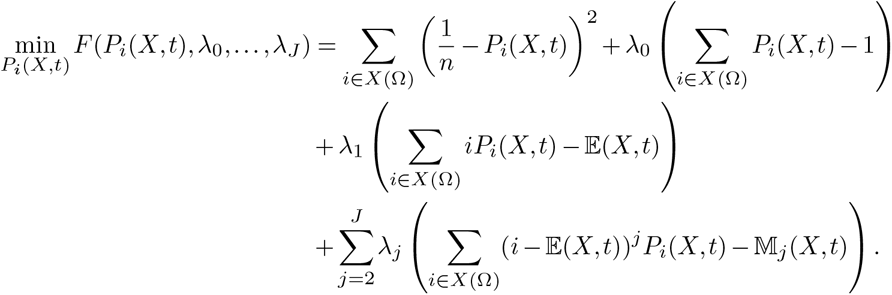

Setting *∂F* (*P* (*X,t*),*λ*)*/∂P*_*i*_(*X,t*) = 0 leads to a solution for *P*_*i*_(*X, t*) in terms of Lagrange parameters, as indicated in the main paper:

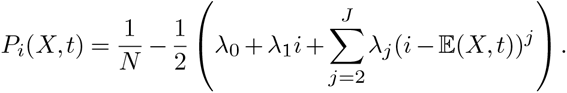

A similar approach has been applied in Smadbeck and Kaznessis (2013). However, it should be emphasized that the inequality *P*_*i*_(*X, t*) is not explicitly considered in this analysis, and hence solutions might violate this condition. This indeed happens for ZIC, as can e.g. be seen in Figure 9A.

## Supplementary Note 3: Plausibility Check

### A. Moment inequalities for the curtosis

Let *X* be a random variable on the finite support *X*(Ω) = {0, 1, …, *n* – 1}. Similar to the derivations in the main document, we bound the curtosis ℂ(*X*) using the following inequality

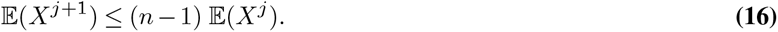

In addition, we rely on two equations that follow directly from the definition of the variance and skewness, respectively:

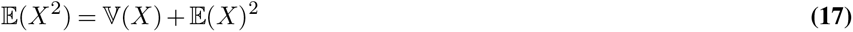

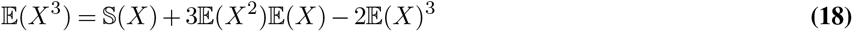

From this we eventually conclude

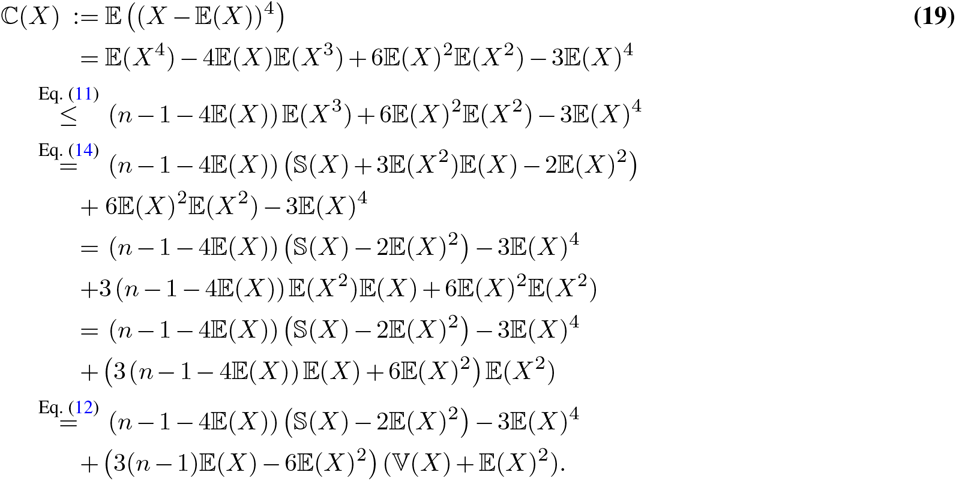

### B. Sharpness of inequalities

Considering a distribution of the type

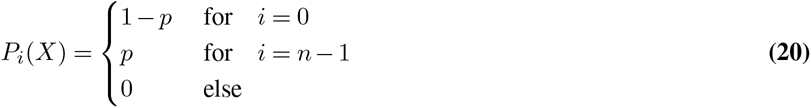

for *p ∈* [0, 1] eveals that the inequalities are sharp. Indeed, all inequalities derived by our approach become equalities when applied to Eq. (20). The prove this, we recapitulate that the only inequality we use during the derivation of our necessary conditions is Relation Eq. (11). For our candidate distribution it holds that

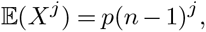

which directly implies Inequality Eq. (11) to be an equality.

## Supplementary Note 4: Simulation Results for Systems 2 and 3

Results for Systems 2 and 3 analogous to those shown in the main manuscript for System 1 are illustrated in Figures 6 and 7 for a maximum of 250 optimization steps, and in Figures 8 and 9 for a maximum of 25 optimization steps, respectively.

**Fig. 6.**
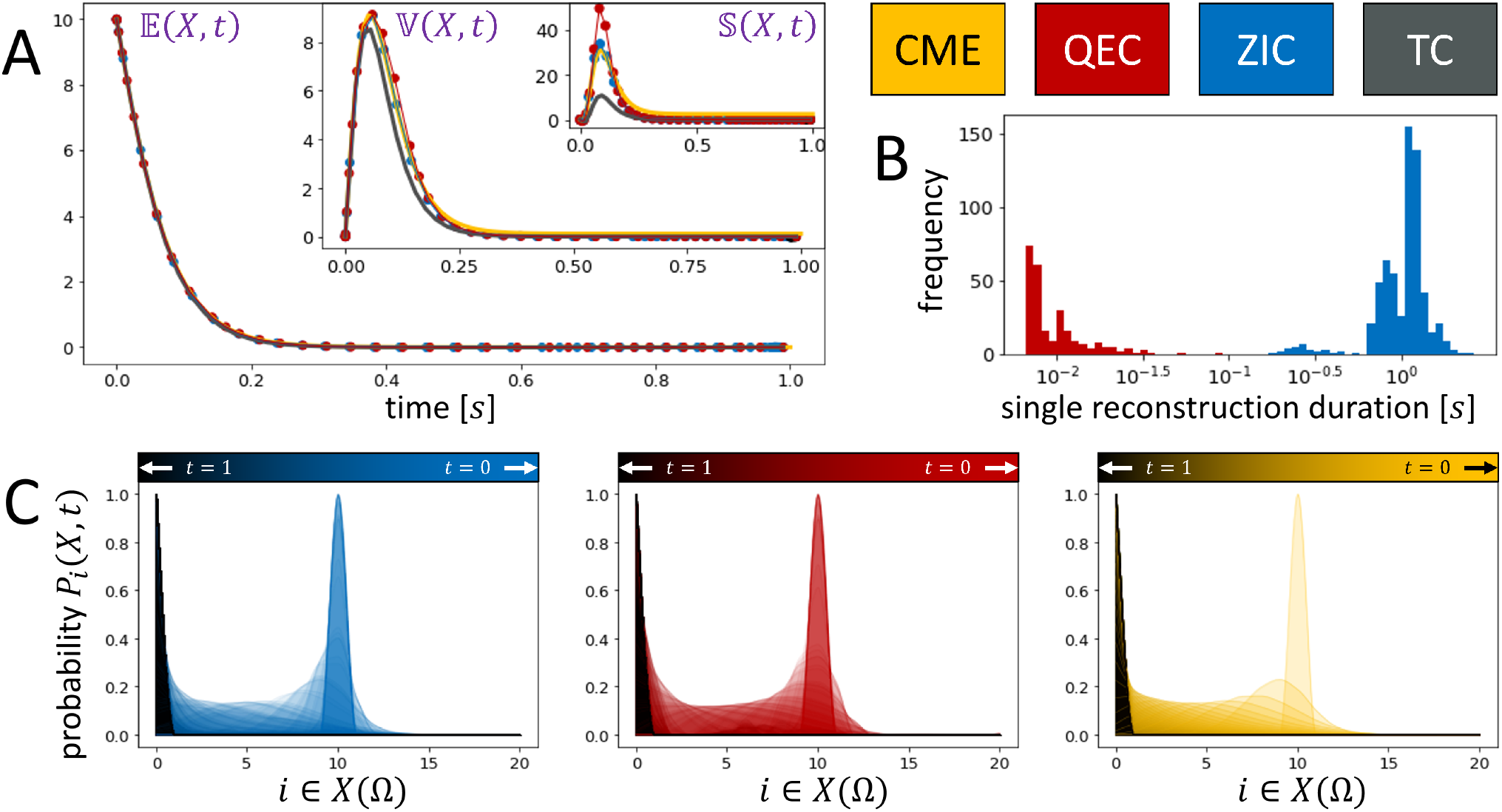
Comparison results for System 2 and 250 optimization steps. Comparison of MoM results for different closure schemes for System 2 with the first three moments and *X*(*t* = 0) = 10 and *n* = 21. Reference moment trajectories and distribution courses obtained from the CME are depicted in yellow. For ZIC and QEC, a maximum of 250 optimization steps was allowed per distribution reconstruction. (A) Courses of 𝔼, 𝕍 and 𝕊 for ZIC (blue), QEC (red) and TC (gray). For ZIC and QEC, each dot on the line corresponds to one distribution reconstruction step. (B) Histogram of computation times for single reconstruction steps. (C) Continuous interpolation of the reconstructed discrete distributions for ZIC and QEC in comparison to the reference course. Light colors represent early time steps, darker colors correspond to reconstructions in later steps.

**Fig. 7.**
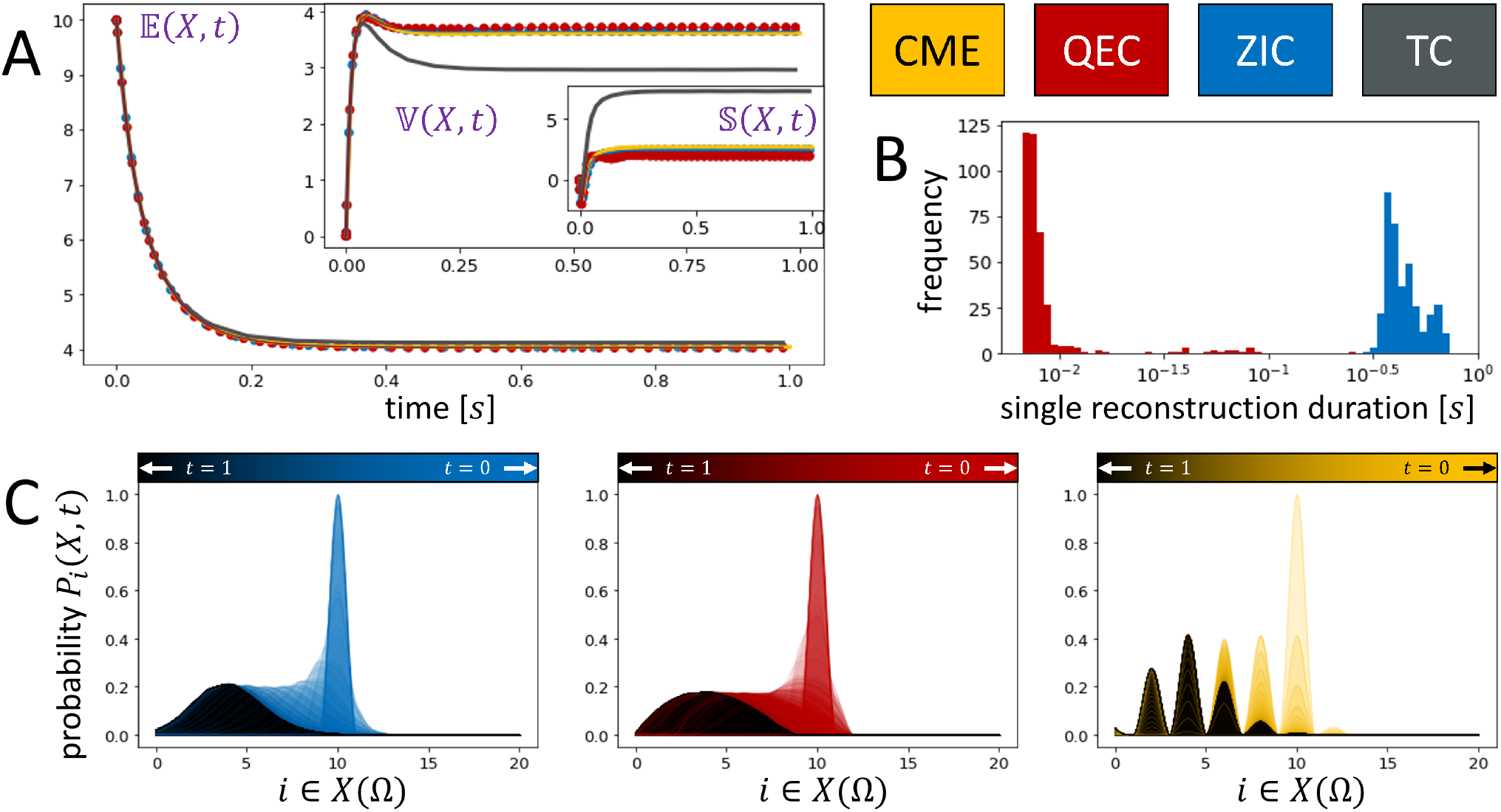
Comparison results for System 3 and 250 optimization steps. Comparison of MoM results for different closure schemes for System 2 with the first three moments and *X*(*t* = 0) = 10 and *n* = 21. Reference moment trajectories and distribution courses obtained from the CME are depicted in yellow. For ZIC and QEC, a maximum of 250 optimization steps was allowed per distribution reconstruction. (A) Courses of 𝔼, 𝕍 and 𝕊 for ZIC (blue), QEC (red) and TC (gray). For ZIC and QEC, each dot on the line corresponds to one distribution reconstruction step. (B) Histogram of computation times for single reconstruction steps. (C) Continuous interpolation of the reconstructed discrete distributions for ZIC and QEC in comparison to the reference course. Light colors represent early time steps, darker colors correspond to reconstructions in later steps.

**Fig. 8.**
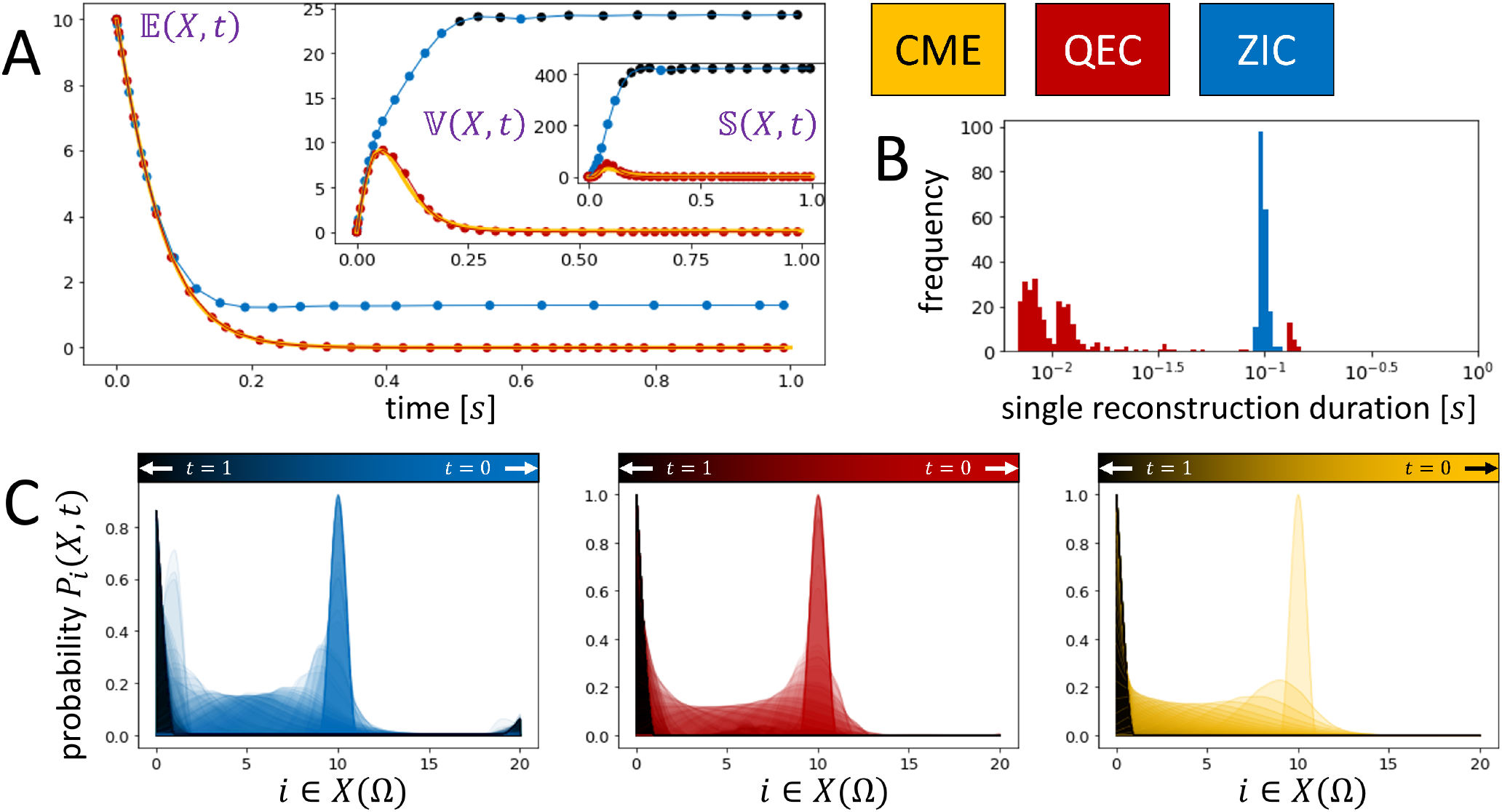
Comparison results for System 2 and 25 optimization steps. Results similar to Figure 6 with the maximal number of optimization steps set to 25 for QEC and ZIC for each reconstruction step.

**Fig. 9.**
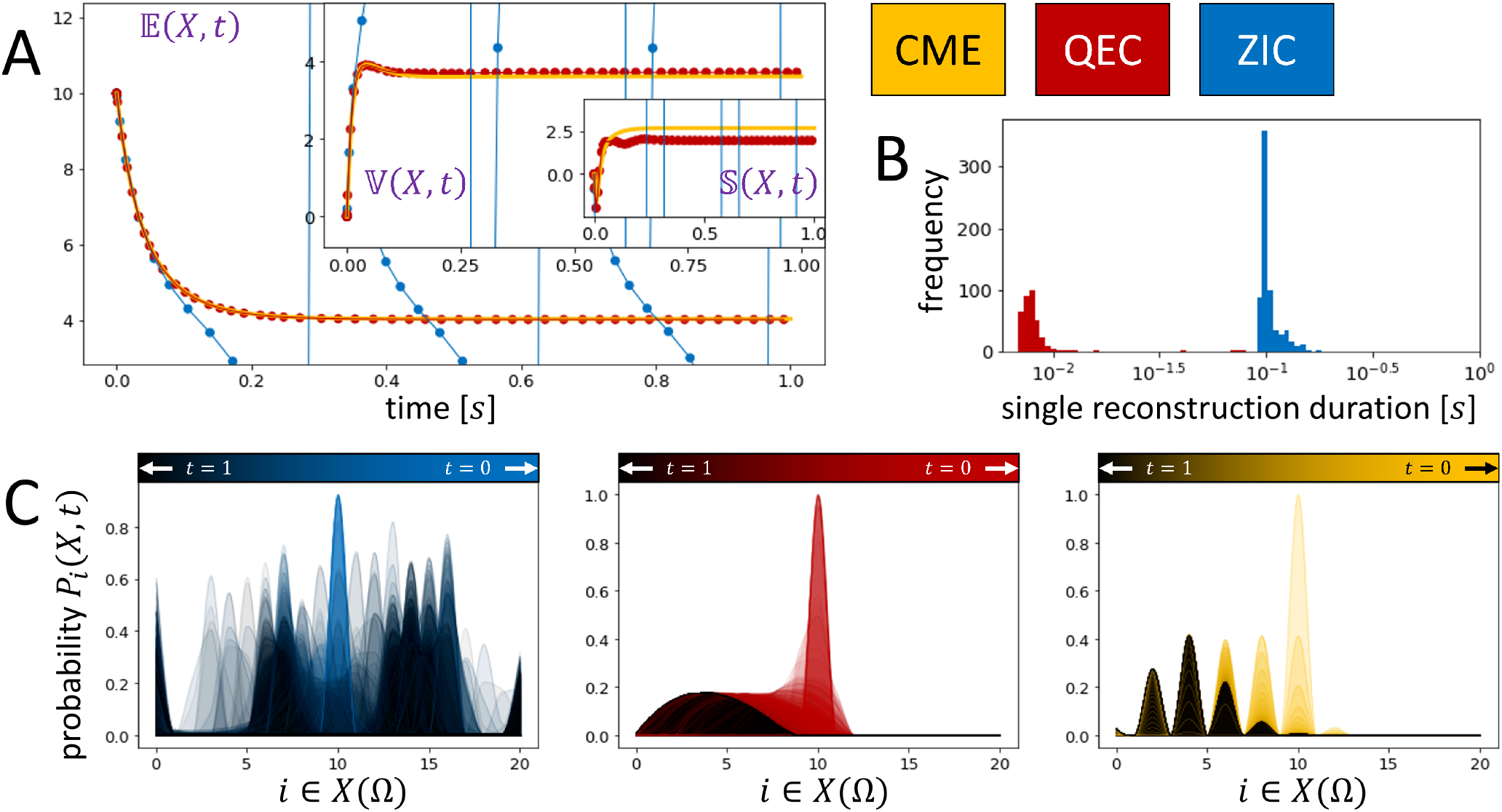
Comparison results for System 3 and 25 optimization steps. Results similar to Figure 7 with the maximal number of optimization steps set to 25 for QEC and ZIC for each reconstruction step.

